# RNAcare: Integrating Clinical Data with Transcriptomic Evidence using Rheumatoid Arthritis as a Case Study

**DOI:** 10.1101/2025.01.31.635914

**Authors:** Mingcan Tang, William Haese-Hill, Fraser Morton, Carl Goodyear, Duncan Porter, Stefan Siebert, Thomas D. Otto

## Abstract

**Background:** Gene expression analysis is a crucial tool for uncovering the biological mechanisms that underlie differences between patient subgroups, offering insights that can inform clinical decisions. However, despite its potential, gene expression analysis remains challenging for clinicians due to the specialised skills required to access, integrate, and analyse large datasets. Existing tools primarily focus on RNA-Seq data analysis, providing user-friendly interfaces but often falling short in several critical areas: they typically do not integrate clinical data, lack support for patient-specific analyses, and offer limited flexibility in exploring relationships between gene expression and clinical outcomes in disease cohorts. Users, including clinicians with a general knowledge of transcriptomics, however, who may have limited programming experience, are increasingly seeking tools that go beyond traditional analysis. To overcome these issues, computational tools must incorporate advanced techniques, such as machine learning, to better understand how gene expression correlates with patient symptoms of interest.

**Results:** Our RNAcare platform, addresses these limitations by offering an interactive and reproducible solution specifically designed for analysing bulk RNA-Seq data from patient samples in a clinical context. This enables researchers to directly integrate gene expression data with clinical features, perform exploratory data analysis, and identify patterns among patients with similar diseases. By enabling users to integrate transcriptomic and clinical data, and customise the target label, the platform facilitates the analysis of the relationships between gene expression and clinical symptoms, like pain and fatigue. This allows users to generate hypotheses and illustrative visualisations/reports to support their research.

As proof of concept, we use RNAcare to link inflammation-related genes to pain and fatigue in rheumatoid arthritis (RA) and detect signatures in the drug response group, confirming previous findings and generating new hypotheses.

**Conclusion:** We present a novel computational platform allowing the interpretation of clinical and transcriptomics data in real-time. The platform can be used for data generated by the user, such as the patient data presented here or using published datasets.

The platform is available at https://rna-care.mvls.gla.ac.uk/, with its source code at https://github.com/sii-scRNA-Seq/RNAcare/.

## BACKGROUND

Autoimmune and autoinflammatory diseases incorporate a heterogeneous group of chronic diseases with substantial morbidity and mortality, thereby posing significant health challenges. In these conditions, immune dysfunction results in chronic inflammation and damage to various organs, including the joints (as seen in rheumatoid arthritis (RA)), the skin (as in psoriasis), and vital internal organs such as the heart and kidneys (as in systemic lupus erythematosus). Over time, autoimmune diseases can severely reduce quality of life, lead to long-term disability, and often require lifelong treatment to manage symptoms and prevent further damage. While there have been substantial advances in the treatment of RA and related rheumatic immune-mediated conditions based on an improved understanding of the underlying disease pathogenesis, there remains a significant unmet clinical need. Treatment responses remain unpredictable and inadequate response is common; some patients may respond to drugs with one mode of action, while others may respond to another drug or not at all. Currently, our ability to stratify (or personalise) treatment such that the right patient receives the right drug for their rheumatic disease is very limited [1]. Furthermore, despite improvement in inflammation levels with treatment, many patients with RA and similar immune conditions report persistent fatigue and pain despite treatment [2, 3]. However, multiple studies have shown discordance between physicians’ assessments of inflammation and patient-reported pain making the understanding of these immune diseases challenging.

One approach to study the underlying disease processes is to use transcriptomic data from relevant patient cohorts and correlate them with different phenotypes, like drug response/resistance, disease severity, disease mechanisms, or patient-reported symptoms, including pain and fatigue. Several studies have indicated that genes, transcripts and proteins associated with pain can be identified [4, 5], supporting the possibility that integrating transcriptomic and clinical data from well-phenotyped disease cohorts may provide further insights into disease mechanisms and patient outcomes. However, most analyses are performed by trained bioinformaticians who have relatively limited experience with the patients or diseases being studied, creating a data-analysis bottleneck, while clinicians often do not have the analytic skills to do this work themselves.

In response to this challenge, several tools have been developed to try and simplify transcriptomic analysis, especially through web-based platforms that streamline setup and configuration for users. The Shiny framework has facilitated the development of web interfaces for R-based pipelines, contributing to the growth of web applications for gene expression analysis `Click or tap here to enter text.`. Several computational tools exist to analyse transcriptomic data and visualise them through web frontends. However, most have limitations, see supplemental Table 1: Phantasus [9], a web platform, integrates a highly interactive JavaScript heatmap interface with an R backend, supporting essential steps in gene expression analysis such as data loading, annotation, normalisation, clustering, differential expression, and pathway analysis. However, Phantasus does not support transcriptomic dataset integration. Similarly, CFViSA [10] provides a comprehensive platform for omics-data visualisation and statistics, integrating microbiome and transcriptome analyses, but lacks support for batch integration and machine learning extensions. AmiCa [11], another web server for analysing microRNA and gene expression in cancer, does not accept user-uploaded data, while GEOexplorer [12] is limited to integration of two datasets, and does not include many options to visualise the batches. Notably, none of the aforementioned tools effectively integrate multiple cohorts of transcriptomic with clinical data.

In response to these limitations and the need to allow easy integration and analysis of transcriptomic datasets aligned with clinical phenotype in rheumatic disease cohorts, we present RNAcare, a web application designed for integrated gene expression and clinical data analysis. We first describe how our tool compares to commonly used existing tools and then report three case studies using RNAcare to perform transcriptomic analysis associated with key clinical parameters (pain, disease activity, fatigue and treatment response) in three RA patient cohorts.

## IMPLEMENTATION

The overall aim of RNAcare is to provide a web-based tool simplifying the integration and analysis of transcriptomics and clinical data showing the analysis workflow in tabs as outlined in Figure 1 and described later in more detail. RNAcare was developed in Python, utilising several open-source packages. The web server was built using the Django framework, adhering to the FAIR (Findable, Accessible, Interoperable, and Reusable) principle [13], see Supplemental Figure 1. The platform employs the Plotly graphics system for generating interactive visualisations on the fly. The platform can be installed locally from https://github.com/sii-scRNA-Seq/RNAcare/.

**Figure 1.**
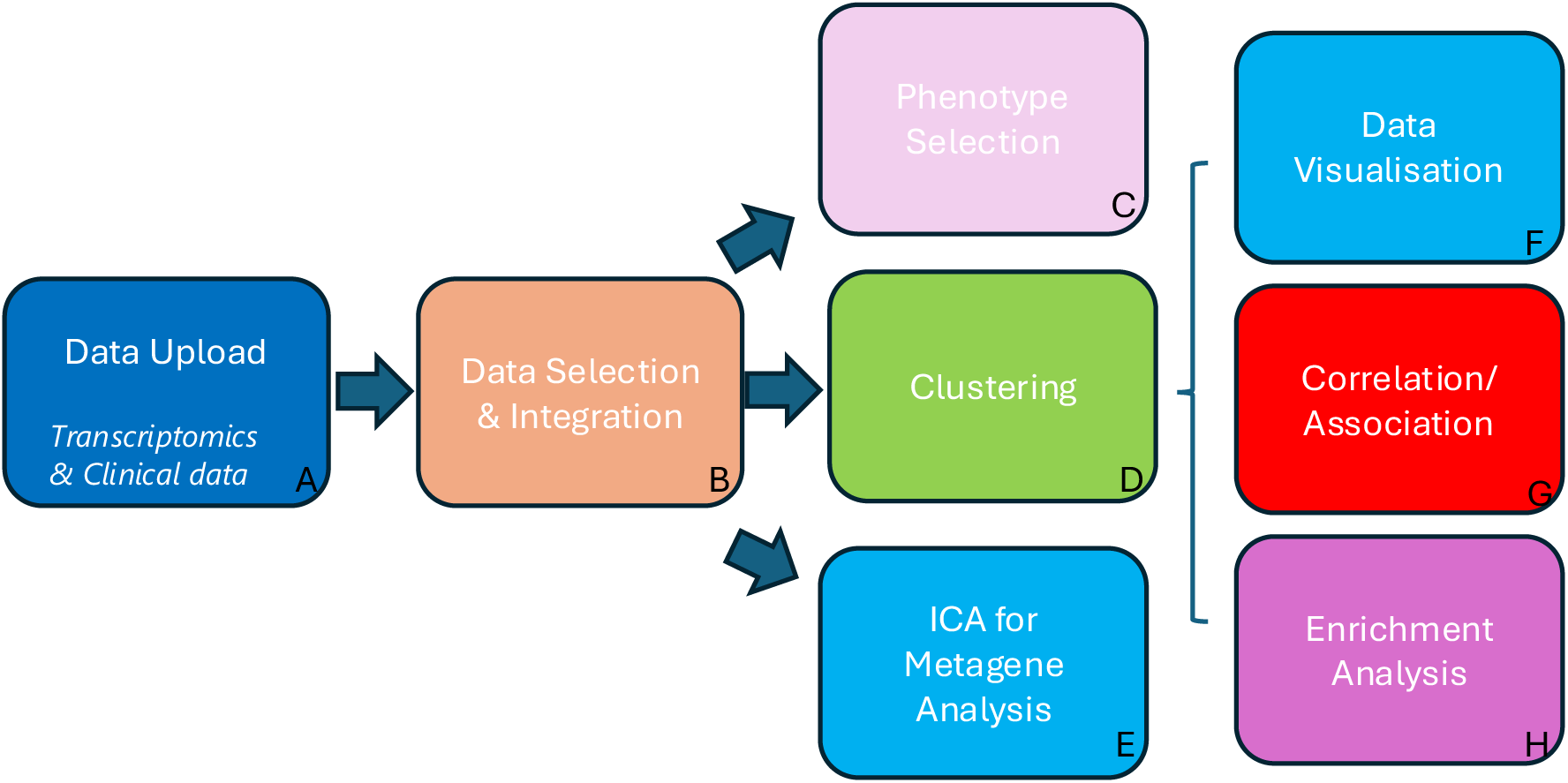
RNAcare workflow overview. (A) In the first tab, users can upload transcriptomics, proteomic and clinical data (optional); For proteomic data, some of the results from module G, H may be not highly reliable based on the current design on transcriptomics. (B) In the data Selection and Integration, users can combine different datasets and have the option to select different integration methods; (C) Phenotype Selection: new labels can be created and added from the clinical data (continuous or discontinuous) for downstream analysis; (D) Clustering: the transcriptomic data is visualised with cluster-based methods as well as projected into 2D; (E) Independent Component Analysis (ICA) for metagene analysis; (F) Data visualisation for labels/clusters; (G) Correlation/Association is to find relationship between transcriptomic data, clinical data and the label using Differential Expression or machine learning; (H) Enrichment Analysis for user’s cluster/label of interest.

To enable several users working on the system, Django is used to allow group and authentication management. System administrators can easily assign group roles to specific users, enabling them roles to only view and operate the permitted data. To improve the performance of the system we used Nginx server caches.

### Datasets

For testing RNAcare we used three RA datasets (Table 1) – Optimal management of RA patients requiring Biologic Therapy (ORBIT), Pathobiology of Early Arthritis Cohort (PEAC) and RA-MAP as part of the IMID-Bio-UK consortium.

**Table 1.** Overview of the PEAC, ORBIT and RA-MAP cohorts used in RNAcare. The clinical features are listed in the Supplemental Table 2.

| Cohort name | Number of participants | Data type | tissue type | #clinical feature |
| --- | --- | --- | --- | --- |
| PEAC [14] | 49 | Bulk RNA-seq | Blood | 7 |
| ORBIT [15] | 139 | Bulk RNA-seq | Blood | 16 |
| RA-MAP [16] | 63 | Relative fluorescence units(RFU) | Plasma | 12 |
|  | 34 | Microarray | Whole Blood |  |

The Pathobiology of Early Arthritis Cohort (PEAC) [14] was established with the aim to create an extensively phenotyped cohort of patients with early inflammatory arthritis, including RA, linked to detailed pathobiological data. The clinical and transcriptomic data can be found on EBI ArrayExpress with accession E-MTAB-6141. ORBIT [15] was a study comparing Rituximab to anti-TNF treatments. Blood was taken from patients before drug treatment. The RA-MAP Consortium [16] is a UK industry-academic collaboration to investigate clinical and biological predictors of disease outcome and treatment response in RA, using deep clinical and multi-omic phenotyping. The raw data were retrieved from accessions GSE97810 and GSE97948 on ArrayExpress. The clinical data can be found here: https://doi.org/10.6084/m9.figshare.c.5491611.v1

#### Datasets for tool comparison

For comparison with other platforms, we used their example datasets. GSE53986[17] for Phantasus, and GSE106382[18] and GSE20589[19] for GEOexplorer.

#### Clinical data used from the cohorts

RA disease activity was assessed using the validated DAS28 (Disease Activity Score using 28 joint counts) score, which was calculated from the original recorded 28 joint counts plus a blood marker of inflammation, typically the erythrocyte sedimentation rate (ESR) or the C-reactive protein (CRP) level [20]; Pain VAS is short for pain Visual Analog Scale (VAS) for self-reported pain, which is a unidimensional measure of general pain intensity, used to measure patients’ current pain level (no pain 0–4 mm; mild pain 5-44 mm; moderate pain 45–74 mm; severe pain 75–100 mm), which can be tracked over time, or used to compare pain severity between patients [21]. The pain VAS is not specific to RA and has been widely used in a range of patient populations, including those with other rheumatic diseases, patients with chronic pain, cancer, or even cases with allergic rhinitis [22]; The fatigue VAS ranging between 0 and 100 cm. Fatigue was considered mild if the fatigue VAS was <20 cm, moderate if 20≤VAS<50 and severe if VAS≥50 cm [23]. All this clinical information is stored on csv files (pain VAS for PEAC and ORBIT and fatigue VAS for RAMAP) and, after curation, loaded into RNAcare.

### A - Data Upload

The user has the option to use data uploaded previously, upload their data on the fly or a combination of both. As proof of concept, we included the three RA datasets described above, including clinical and expression data. RNA-Seq requires a read count matrix, which will be normalised later. Other omics data need to be normalised initially before uploading. Clinical data are uploaded as a table.

### B - Data Selection and Integration

Before data integration, RNAcare detects whether the format of the expression data are integers or non-integers. For integers, the platform handles the expression data as RNA-Seq data (applies to PEAC and ORBIT), which are transformed from raw counts to CPM (count per million). For non-integers, for example microarray/proteomic data, the platform handles the expression data as normalised data (applies to RAMAP), so users need to pre-normalize these data types.

The user has the option to log1p transform their data before harmonisation. RNA-Seq data is often highly skewed, with large differences in scale between genes. Log transformation helps stabilise the variance and makes the data more suitable for batch correction. If this option is selected, both the gene expression data and clinical numeric data will undergo log1p transformation. Conversely, if the log1p transformation is not selected, the gene expression data will remain unchanged, while clinical numeric data will be normalised to a 0–1 range.

For our case studies, we obtained the best integration results with the log1p normalisation. Therefore, for Case Study 1, we used log1p to integrate PEAC and ORBIT data; for Case Study 2, we used log1p to process the proteomics data but did not log1p transform the microarray data (because it has already been processed in the microarray data); for Case Study 3, we used log1p to transform ORBIT data.

Next, the user as two options for the integration, Harmony [24] and Combat [25], and three options for feature reduction, principal component analysis (PCA) [26], t-distributed stochastic neighbour embedding (t-SNE) [27] and Uniform Manifold Approximation and Projection (UMAP) [28]. In general, it is unclear a priori which integration will generate the best data, and this will vary for different datasets, the user has different options for this in RNAcare.

### C - Phenotype Selection

Users can define clinical classes, categorise continuous clinical parameters and generate new user-defined labels based on numerical variables from uploaded meta files (categorical variables in the meta files can be redefined for labelling). This feature allows for the creation of customised fields that can be used as targeted dependent variables in subsequent analysis.

### D - Clustering

Clustering allows grouping of patients by common expression patterns to facilitate the comparison of gene expression between different sample groups in order to identify differentially expressed genes (DEGs). We implemented various methods including K-means [29], Leiden [30], and HDBSCAN [31] to allow the user to choose their method of choice.

The default parameters, for K-means, is the number K, which needs to be set before running the clustering. The process will be terminated if users use a very large K or the size of any of the clusters in the sample is less than 5. For Leiden, resolution is set to 1 by default. For HDBSCAN, the default parameter is minSize, representing the minimum size of clusters with a default value of 5 set in the interface.

Users can easily trace the clustering procedure by specifying the desired range and number of levels. The algorithm will automatically generate clusters based on the intervals within the specified range.

### E - ICA for Metagene Analysis

Independent component analysis (ICA) attempts to decompose a multivariate signal into independent non-Gaussian signals. It has already been proved its successful application in computational biology [32]. We introduced ICA into transcriptomics to decompose the gene expression matrix into several independent components. Each independent component was treated as a metagene made up of explainable genes, characterised by a co-expression pattern and was associated with certain meaningful biological pathway. In practice, we suggest that all clinical fields uploaded before must be named as strings starting with “c_”, such as “c_das”, which will be easy for the program to recognise the expression data and apply ICA algorithm.

### F - Data Visualisation

Users can plot genes of interest based on clusters defined in earlier steps, with options to create violin plots, density plots, and heatmaps. The system can also suggest candidate genes through predefined algorithms.

### G - Correlation/Association

Here, the user has the option to associate selected clinical data parameters with the transcriptomic data in order to study phenotypes. Users can select the top N (default = 4) differential genes in each group after clustering. The platform can also apply the Lasso [33] and Ridge [34] method to demonstrate feature importance. The default parameters for regularisation (Lasso and Ridge) are cv=5 for 5 folds cross validation, solver=”saga”, class_weight=”balanced” for potential imbalanced issues. Before regularisation, data needs to be standard scaled for either algorithm to compare all features equally. When we analyse the relationship between signatures and phenotype, Lasso is applied out of the consideration of feature reduction; When we analyse the relationship between metagenes and phenotype, Ridge is applied.

### H - Enrichment Analysis

As the previous analysis will return gene lists of interest, this tab helps to identify biological pathways and gene sets significantly enriched in the dataset. Users can perform GO enrichment [35] analysis for specific clusters, enhancing their understanding of the underlying biological processes.

Another feature of RNAcare is download all the results of the different analysis steps, results as csv and figure as SVG for further processing and use in publications.

## Results

Here we present the first tool that allows users without bioinformatics skill to integrate clinical and transcriptomics data. Although there are similar tools available (Supplemental Table 1), none has all the features of RNAcare. First, we compared it with two other tools to show that it is comparable to other tools. Further, we show RNAcare has more features in terms of data integration, contributor visualisation and DEGs for clustering, before showing three use cases.

### Comparison with selected existing tools

#### Phantasus

Phantasus is an analysis tool accessible via both web and local applications, designed to analyse transcriptome data derived from microarray or RNA-Seq technologies[9]. It does not allow the use of clinical data.

One dataset used in Phantasus, consisted of 16 samples of bone marrow-derived macrophages, untreated and treated with three stimuli: lipopolysaccharide (LPS), interferon-gamma (IFNg), and combined LPS+IFNg (GSE53986). Both tools, Phantasus and RNAcare, have the option to perform differential gene expression and perform GO Enrichment. The results (Figure 2 A, B) are very similar: RNAcare has a similar GO result.

**Figure 2.**
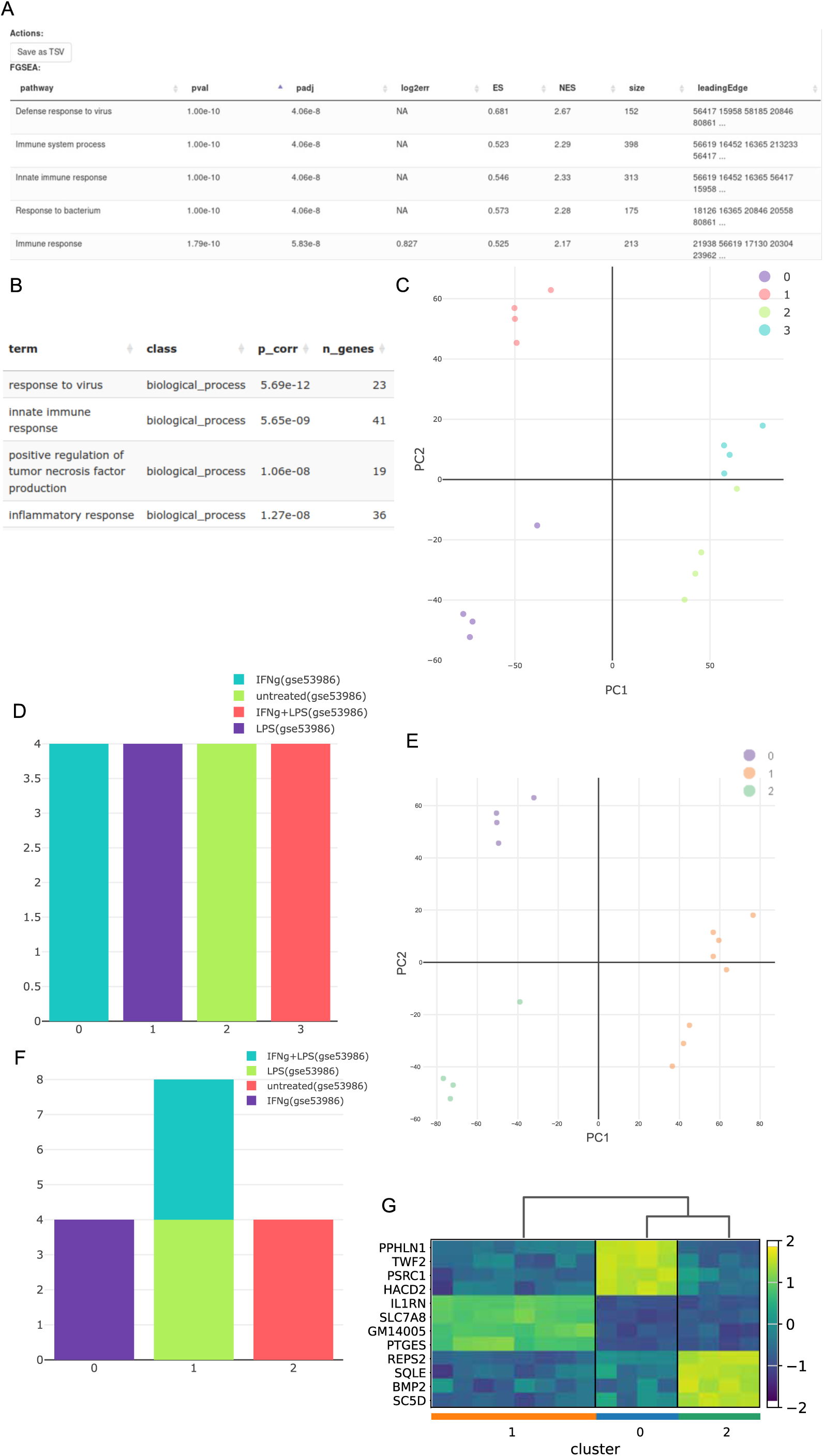
Comparison with Phantasus using GSE53986. (A) Go enrichment of LPS vs untreated with Phantasus. (B) Go enrichment of LPS vs all groups with RNAcare. (C) Visualisation for K means (K=4) clustering using RNAcare while Phantasus doesn’t support. (D) Contributions to the 4 clusters from each label. (E) Visualisation for K means (K=3) clustering using RNAcare (F) Contributions to the 3 clusters from each label. (G) DEG heatmap plot for clusters with RNAcare.

The reason why Phantasus’s p adjusted value is so significant is because it only includes the top 12000 expressed genes and did a binary DE between LPS and untreated group. In RNAcare, we used all genes for analysis, which users can filter before uploading. Besides, we used global DEGs among all the four groups.

However, one useful feature of RNAcare missing in Phantasus is to visualise the clustering of the data. If we use K=4 for clustering, we can see the data can be well clustered into 4 clusters matching with labels. It can be seen that the cluster are separated in the PCA and its visualisation as PCA plot (Figure 2C). For each cluster we plot its contributors by Labels (Figure 2D). We can also see IFNg+LPS and LPS are closed, so if we choose K=3, we will see these two groups merge and find the common signatures compared to the other groups (Figure 2E-G), as cluster 0 has samples from IFNg, cluster 1 samples from LPS and IFNg+LPS and cluster 2 primarily samples from the untreated. NexThe visualisation methods allow users to have a power view of the data. These cluster visualisation methods are not part of Phantasus and are advantageous for RNAcare.

#### GEOexplorer

Next, we compared GEOexplorer with RNAcare. We used two datasets for integration: 1. GSE106382, a dataset of induced pluripotent stem cells (iPSCs) from healthy controls, sporadic amyotrophic lateral sclerosis (ALS) and familial ALS patients including a subgroup of familial cases who carried a pathogenic mutation in the SOD1 gene (SOD1 ALS). We focussed on the SOD1 ALS samples and therefore will subsequently refer to these as model SOD1 spinal motor neurons. 2. GSE20589, which includes cervical spinal motor neurons from healthy controls and SOD1-related ALS post-mortem. These data are microarray data from the GPL570 platform.

Both tools generated allow similar analysis such as differential expression and GO enrichment and basic visualisations of the results (Supplemental Figure 2A,B). However, GEOexplorer only supports integration of two datasets. In addition to allowing integration of multiple datasets, RNAcare also offers visualisation by batches to assess the quality of dataset merging after batch correction, making it well-suited for handling large-scale data integration. In contrast, GEOexplorer reviews batch-corrected results on a record-by-record basis (Supplemental Figure 2C). Again, GEOexplorer does not have the function integrate clinical data.

Overall, RNAcare performs similar as existing tools to standard analysis, however it has a new feature and aninnovation of allowing the user to integrate clinical data. In the use cases we will introduce these features, however we want to highlight the standard of implementation of RNAcare.

#### Computational advantages

Of note, RNAcare uses more modern implementation, namely the RESTful Web Service architecture (Supplemental Figure 3), which provides a flexible and standardised programmatic interface. This architecture facilitates seamless execution of stress tests, enabling the systematic evaluation of server performance under varying load conditions. In comparison, GEOexplorer and Phantasus are Shiny web apps, which inherently limits the programmability and scalability.

Although we have not tested this in other software, RNAcare mitigates potential server abuse, with the evaluated system incorporating a timeout mechanism of 180 seconds for machine learning tasks, ensuring that resource-intensive or malicious requests are automatically terminated upon exceeding the time limit. We stress tested RNAcare with several users accessing it simultaneously, without any issues, in terms of processing and waiting times. For example, integrating and analysing two datasets (PEAC and ORBIT) on our server with 62GB of RAM and 12 threads we observed no delays for up to 20 users, showing the modern implementation of our system.

We next proceeded to test cases using RNAcare to associate clinical parameters of interest with combined transcriptional data in the uploaded RA cohorts.

**Case study 1**: Evaluation of the peripheral RNA signatures associated with pain in RA

Pain is an important clinical symptom in RA which often persists, even when inflammation appears improved with immunomodulatory therapies, suggesting that pain in RA involves more than just inflammation. Of note, Hall et al. [5] identified 128 DEGs associated with inflammation-induced pain in dorsal root ganglia (DRG) samples from five RA patients that were not present in non-arthritic controls RNA-Seq without RA; Youssef et al. [36] demonstrated that S100A9 is overexpressed in the lining layer of inflamed synovial tissue in RA, while Foell et al. [37] highlighted S100A12 was strongly expressed in inflamed synovial tissue whereas it was nearly undetectable in synovia of controls or patients after successful treatment. Serum levels of S100A12 correlated with disease activity.

In this case study, we used two existing RA datasets (ORBIT and PEAC, baseline samples) which are pre-loaded in RNAcare, to find peripheral RNA markers associated with different levels of pain, measured using patient self-reported pain VAS[21]. The aim was to integrate the two datasets, perform different analyses to find the association of pain with different variables, and finally see if existing signatures could be in our datasets.

After importing the clinical and RNA-Seq data from ORBIT and PEAC into the RNAcare platform, we created a new categorical label to define two levels of pain, low and high. The pain VAS variable ranges from 0 to 6 (Supplemental Figure 4). We set a threshold of 4 (log1p(45) ≈ 4), which divided the participants into two groups: no/low pain (low-53) and moderate/high pain (high-135). The advantage of RNAcare is that this label can now be used to perform further analysis, while all other derived labels and the original continuous pain feature were excluded from independent features in the subsequent modelling steps.

First, we integrated the data to account for batch effects. We applied the Combat algorithm [38] which integrated well the two datasets, which can be seen in the next tab of RNAcare, with no dataset clustering on its own (Figure 3A).

**Figure 3.**
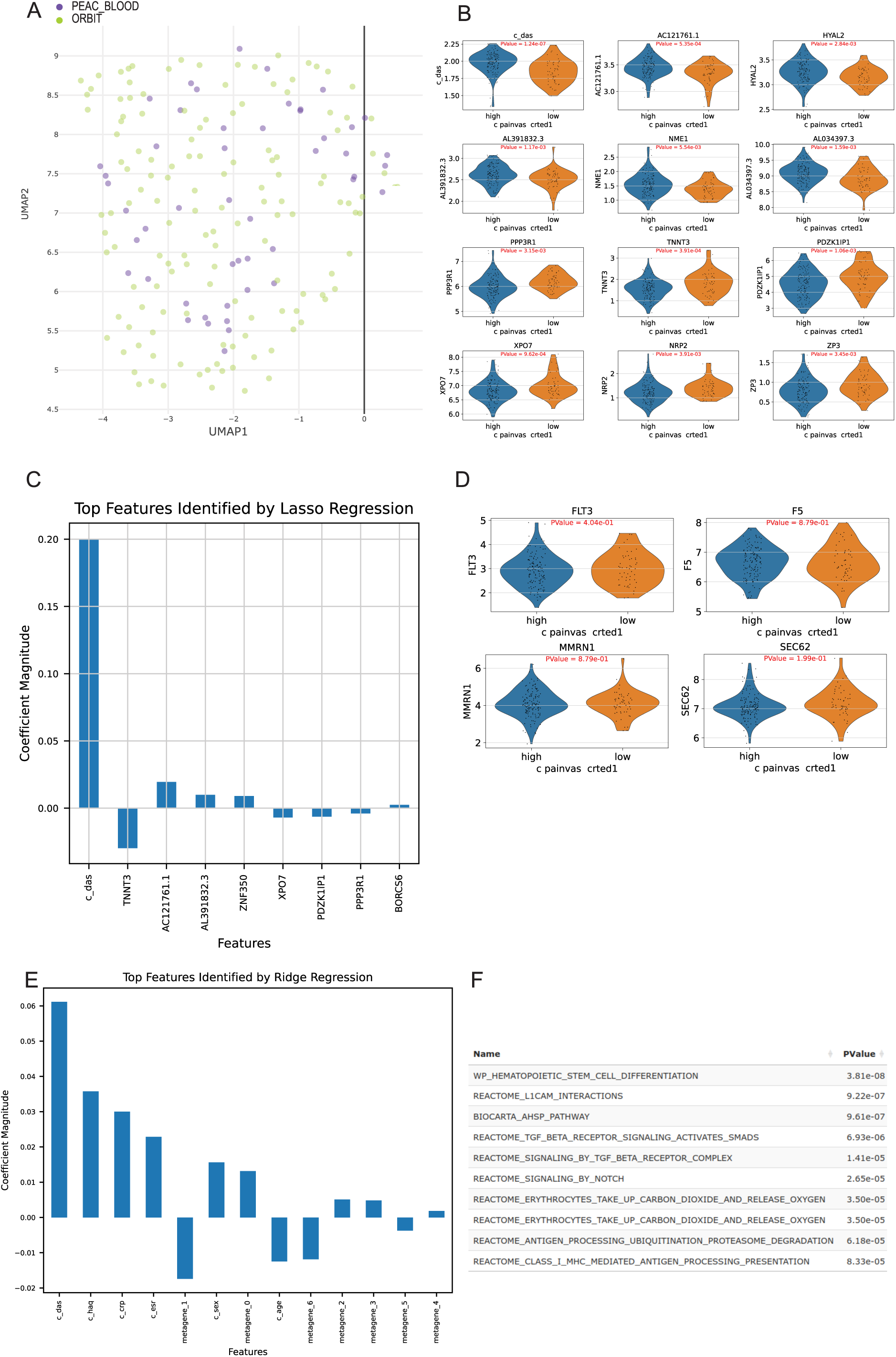
Data Integration.Pain Exploration. (A) Data integration settings and creation of customised labels visualised by UMAP after batch correction (B) Violin plot for DE genes of the labelled groups. (C) Lasso regression with cross validation for high pain group, significant genes are found for the created label. (D) Using RNAcare to visualise the previously identified markers from Bradford et al. (E) Ridge regression with cross validation for high pain group using ICA; (F) Pathway analysis for metagene_1.

This successful integration allowed us to compare the features (genes expression and clinical data) between patients from low and high-pain RA groups. RNAcare uses Scanpy[39] in Python, which expects clinical features as genes. One difference compared to tools like DESeq2 is that Scanpy cannot regress out clinical features when doing DE analysis and DESeq2 assumes data following the negative binominal distribution. During data processing, the platform runs comparison automatically, finding the markers (downloadable from EDA tab) as default candidates when the user wants to plot between different clusters (from DGEA-Target Gene tab). In this case, we find that c_das is significant based on p adjusted value < 0.05 after the comparison. However, Figure 3B still shows the top 12 biomarkers ordered by significance.

To take better advantage of the clinical data, we performed a Lasso regression to identify genes and clinical data associated with the different pain levels. This approach with the parameter class_weight=’balanced’ gives more weight to the patient group with a small number of participants while selecting only the most relevant and predictive features, ignoring irrelevant ones. The coefficient magnitude of every marker after Lasso regression with cross-validation for the high-pain group is plotted in Figure 3C.

Comparing the results of the two methods, in the first comparison only demonstrates DAS score as a significant result and the Lasso regression has a good overlap with other markers, though the order is different. The advantage of the Lasso integration is how it explains the signal. For example, DAS score has the largest co-efficient (0.2), which contributes the most to the pain as expected. This implies that increasing the log-transformed DAS score by 1 unit will result in a exp(0.2)-1=22.14% increase in the odds for patients to be in the high pain group, while controlling the other independent variables unchanged. A negative association is with TNNT3. TNNT3 encodes fast skeletal troponin T protein, and its GO annotation is actin binding. Liu et al. [40] reported that actin binding is a pivotal process in neuropathic pain by regulating spine morphology and synaptic function. Two positively associated transcripts are AC121761.1, a gene associated with expression in the brain[40] and AL391832.3, a long non-coding RNA.

Another feature of RNAcare is to check the expression of existing gene signatures in datasets. In order to see if the DRG biomarkers identified by Hall et al [5] were associated with pain in serum RNA we used RNAcare to visualise these biomarkers. As shown in Figure 3D, those biomarkers show no significant difference in our datasets which are from PBMC; Therefore, they might not be good biomarkers in blood.

In Figure 3E, we used ICA, decomposing the expression data into 7 metagenes. In this analysis each metagene should represent biologically meaningful gene expression pattern of several genes that is statistically independent from other components. Next, we ran Ridge regression of those metagenes with the clinical features over the two pain groups. We can see metagene_1 is mostly highly associated to high pain. Finally, we performed a pathway analysis for the metagene_1 (Figure 3F). Several significant terms returned, including L1CAM interactions and Notch signalling pathways, which has already been proved to be associated with pain in the tissue of DRG, particularly in the context of neuropathic pain [41–43].

Another way to visualise the data in RNAcare is clustering-based methods. To analyse transcripts correlated to disease activity, which are indirectly related to pain, we performed clustering of the two cohorts into two groups using KMeans with two clusters (Figure 4A). The 3200 downregulated and 286 upregulated significant markers (adj Pvalue < 0.05 and |logFC| > 0.5) are identified from cluster 1 (supplemental data file 1, as downloaded from RNAcare, unfiltered), revealing distinct functional roles. We did Gene Set Enrichment Analysis (GSEA) for the clusters (Figure 4B): cluster 1 contains genes upregulated that are involved in inflammation and immune modulation. Notably, transcripts linked to the inflammatory response, such as S100A9, S100A12, and IL1R2, are especially interesting.

**Figure 4.**
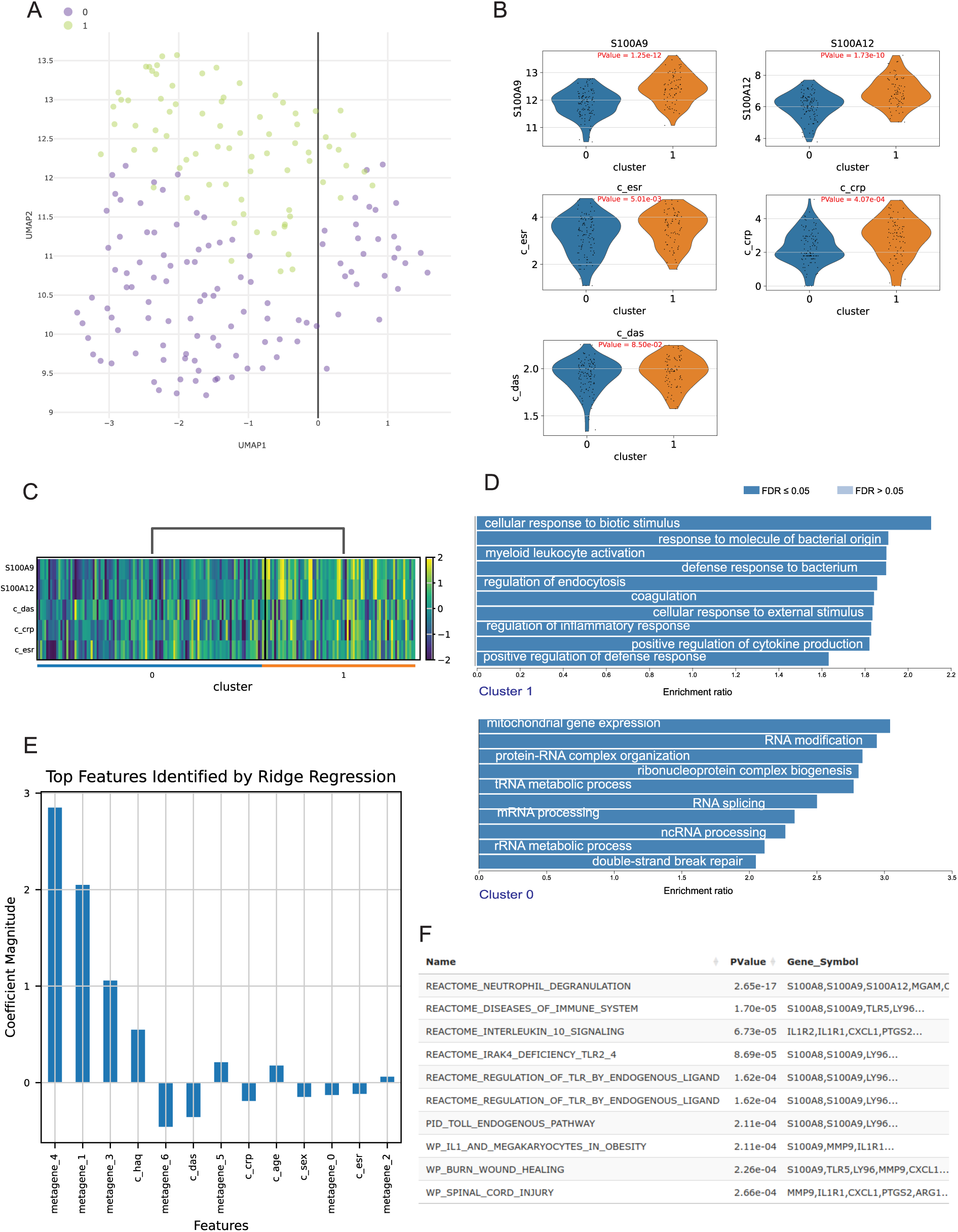
Exploring the clustered data. (A) Clustering the integrated data. (B) GSEA for the clusters. (C) Violin plot for the clusters. (D) Heatmap plot for the clusters. (E) Top features identified by Ridge regression after applying ICA for cluster 1. (F) Pathway analysis for metagene_4.

Additionally, violin and heatmap plots of S100A9, S100A12, disease activity (DAS), Class Switch Recombination (CSR), and Erythrocyte Sedimentation Rate (ESR) showed differences between the two clusters (Figure 4C, D). These findings align with the previous literature mentioned above.

To explore further the clinical features of cluster 1, we used original integrated data, processed it by ICA and ran Ridge regression on cluster 1 (Figure 4E). RNAcare enabled us to perform pathway analysis of individual metagenes, in this case the most significant one metagene_4 (Figure 4F). The top genes were S100A8, ILR1 and again S100A9, which is key of most of the terms.

Overall, we have shown in this case study that it is possible to use the standard pipeline of RNAcare with the pre-loaded data in RNAcare to perform different analyses and generate new hypotheses. Although not many genes were different between the groups, and we could not confirm all the existing data, we can replicate existing results. The difference in the results might come down due to the different cohort. Anyhow, we have shown that it is possible to concatenate analysis, for example combining ICA wit clustering and pathway analysis, highlighting the functionalities of RNAcare.

**Case study 2:** Evaluating the peripheral RNA signatures associated with fatigue in RA

Fatigue is a significant cause of poor quality of life in patients with RA [44, 45] and together with pain, is reported by patients as their most troublesome symptom [46]. The underlying pathophysiology of fatigue in RA, and other chronic immune conditions, remains incompletely understood. Tanaka et al [47] proposed four key mechanistic themes for fatigue in RA, namely inflammation, hypothalamic-pituitary-adrenal axis, dysautonomia, and monoamines, which have a complex network of interconnections between themes, suggesting a key role for inflammatory cytokines in the development and persistence of fatigue.

In this use case, we hypothesise that there are peripheral markers associated with fatigue severity. Similar to the first case, we first tried to find significant biomarkers associated with fatigue. However, we will do that first at the proteomic level and then test the significance of the biomarkers at the transcriptomic level. This sequential workflow allowed us to validate findings and evaluate whether the fatigue-associated signatures were conserved across the proteomic and transcriptomic layers. As for the dataset, we are using the RAMAP cohort, only which included fatigue VAS.

A customised label was created based on fatigue levels (threshold=log1p(50)≈4), stratifying the proteomic samples into two distinct groups (Supplementary Figure 6B). Hence, there are 34 and 29 samples separately in the two groups (Figure 5A). However, when looking for significant differences between the groups, we didn’t find any significant markers whose adjust p values < 0.05. Therefore, we compare it with the result of Lasso and analyse the overlapping 8 markers (Figure 5 B): FGF18, XRCC6, TBP, NGF, KLK3, EIF4A3, USP25, CDC2.

**Figure 5.**
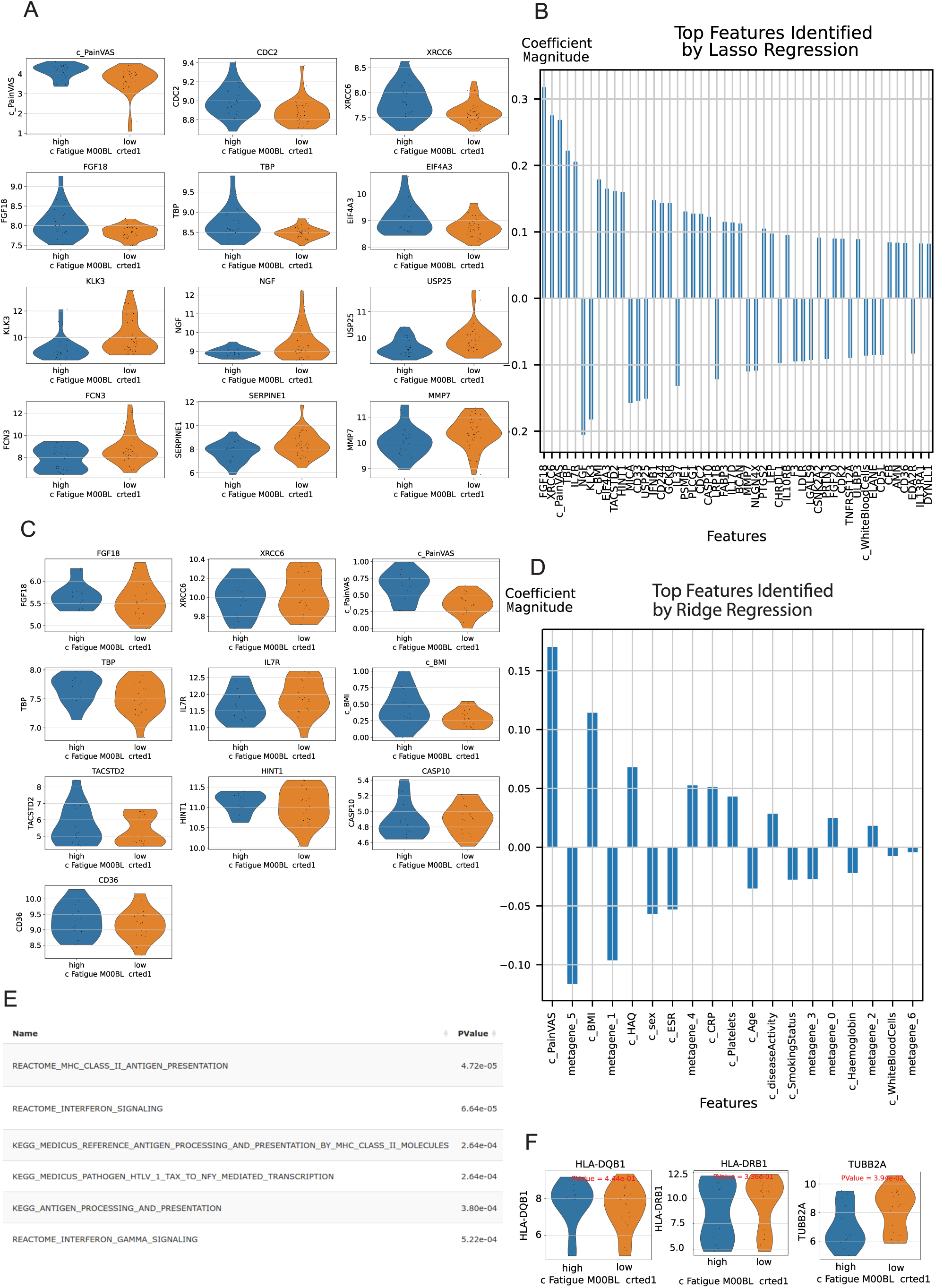
(A) Violin plot for features of interest in different groups to Fatigue. (B) Lasso regression with cross validation to prioritse important features for the group with high fatigue scores. (C) Violin plot for corresponding features at transcriptomic level after running Lasso on Proteomics data. (D) Ridge regression with cross validation for metagenes combined with clinical features. (E) Pathway analysis for metagene_5. (F) Violin plot for genes in metagene_5.

To confirm the Lasso results, we uploaded the microarray data with the RA-MAP cohort, which yielded 34 records (14 high fatigue, 20 low fatigue, Figure 5C). Overall, there is a clear trend confirming the Lasso result. However, none of the results were statistically significant.

Finally, we read around evidence of those found hits and found potential involvement in brain disease [46, 48–50]; we felt this was cherry-picking and would argue that larger sample sizes are needed to obtain more confident results. More important for us is the fact that now the pipeline exists to generate those hypotheses once more data are available.

To explore the data further anyhow, in Figure 5D, we used ICA, decomposing the expression data into seven metagenes by experience and ran Ridge regression with clinical features for high fatigue group. We can see metagene_5 is the most important in the metagenes and we did pathway analysis for it in Figure 5E. We can see its significance in interferon gamma signalling, which is negatively associated with fatigue for chronic/inflammatory diseases, as previously reported [51, 52], and TUBB2A in the metagene might be a potential biomarker involved [51]. The lower expression of TUBB2A in RA is also proved by [52]. In Figure 5F, we see in the high fatigue group, since TUBB2A is lower, and HLA-DQB1/HLA-DRB1 are unchanged, this can lead to the hyperactivation and exhaustion of T cells, which will lead to a decrease in IFN gamma production [53–55]. We also find that taking Lactobacillus acidophilus may potentially increase the level of IFN gamma to relieve fatigue in the inflammatory scenario [56, 57].

**Case study 3:** Evaluating peripheral baseline DEGs associated with treatment response

Resistance to drug treatment is a major problem in treatment responses of RA [58]. Approximately 30%-40% of patients fail to respond to anti-TNF treatment. Although treatment response is likely to be multifactorial, some markers have been replicated consistently between studies [59].

To highlight the functionality of RNAcare, we used a subset of the ORBIT dataset, focusing on exploring the potential genes at baseline associated with responders and non-responders **only** for Anti-TNF after 6 months of treatment. We grouped patients with remission and good response as the responder group (N=35) and patients with partial/non-response as the non-responder group (N=6) by using the metadata provided.

We performed the same pipeline as before, generating a differential expression (DE) of the data with Scanpy and the association with Lasso. After DE (Figure 6A), we got two significantly upregulated DEGs: C19orf53, SLC19A2 and two downregulated DEGs: CTXN2, AC021483.1 for the non-responder group. After the intersection with the Lasso results (Figure 6B), we were left with C19orf53, found in IFN cDNA libraries, producing the protein L10, an interferon-stimulated gene (ISG) product still not well characterised [60]..

**Figure 6.**
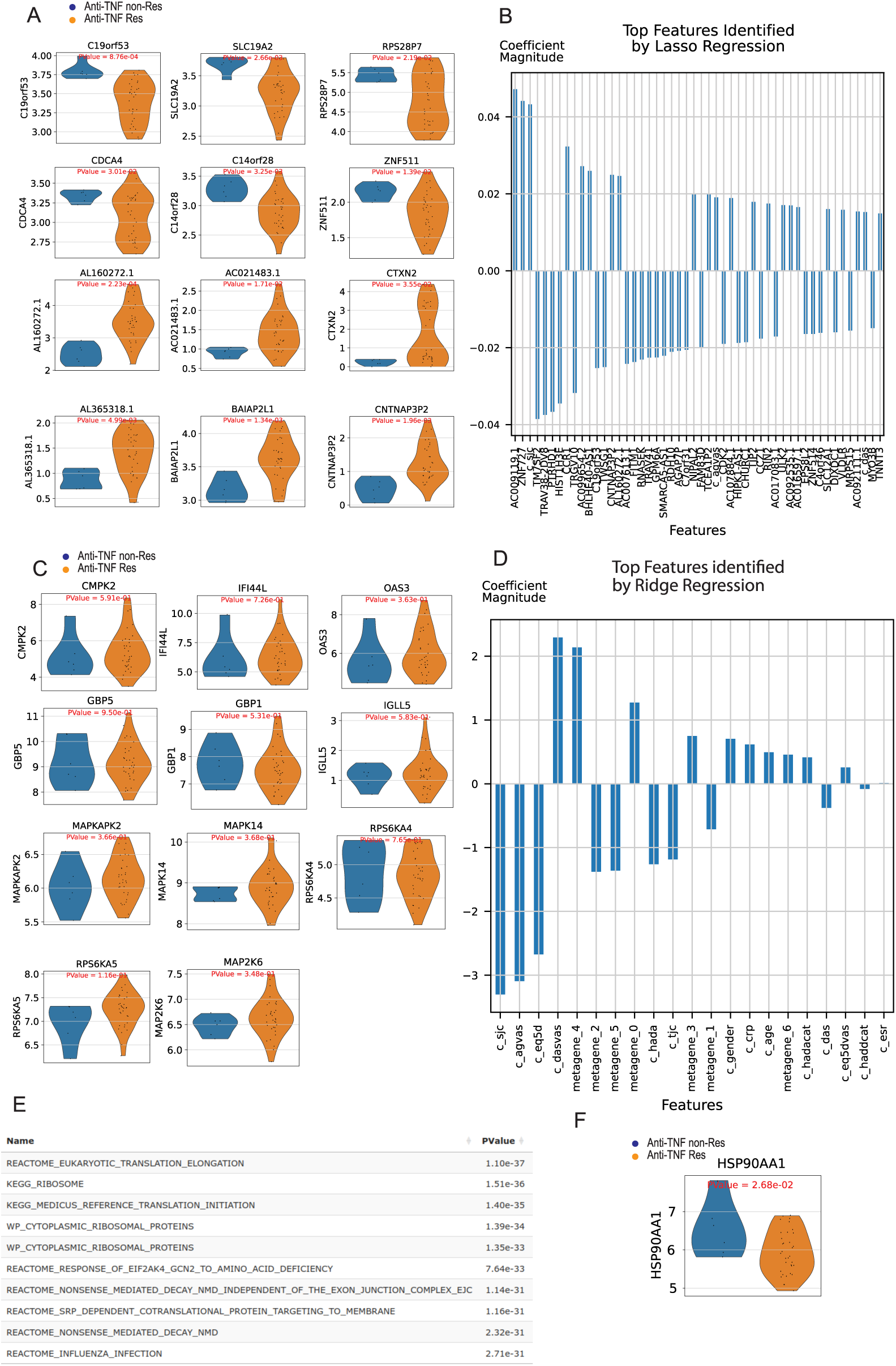
Treatment Comparison. (A) Violin plot of the two groups using Differential Expression. (B) Lasso regression with cross validation for response group. (C) Gene verification for other literatures. (D) Ridge regression with cross validation for response group with metagenes. (E) Pathway analysis for metagene_4. (F) Expression of gene HSP90AA1 from metagene_4 between two groups.

Previously, Yu et al. [61] showed that 41 response-associated genes in 82 RA patients shared common immune pathways including type I IFN signalling, and they reported 6 upregulated DEGs, namely CMPK2, IFI44L, OAS3, GBP5, GBP1, IGLL5, for non-response group and 305 DEGs for response and moderate response groups; Thurlings et al. [62] showed that there was a better clinical response to rituximab in the IFN low signature group; Coulthard et al. [63] reported associated markers at proteomic level, including MAPKAPK2, MAPK14, RPS6KA4, RPS6KA5 and MAP2K6.

We took those gene lists and looked for differences in our datasets. Although we detected those transcripts (Figure 6C), none of the results were statistically significantly different between the two groups. Again, maybe because our sample size was very small, we could not find significant results.

Finally, in Figure 6D, we used ICA, decomposing the expression data into 7 metagenes and ran Ridge regression with clinical features for non-response group. We can see metagene_4 is the most important in the metagenes. It contains the genes HSP90AA1, many RPL and RPS. We performed a pathway analysis (Figure 6E). Among all significant pathways, we find influenza infection, which is highly related to type I IFN signalling pathway [64]. One gene HSP90AA1 (Heat Shock Protein 90 Alpha Family Class A Member 1, Figure 6F), was associated with drug resistance [65]. However, due to the strong ribosome genes this signal could also just be stress. Although we found no strong results, we have proposed another analysis pipeline for the field, which can be easily repeated on our web server.

## CONCLUSION

RNAcare has been designed to enable gene expression analysis of user uploaded/built-in gene expression data with clinical data without the need to be proficient at programming or have advanced bioinformatics skills. One of the downsides of - web-based tools to generate hypotheses from published data is that the tools cannot keep track of the number of experiments performed to correct for multiple testing. Therefore, the user might not be to generate a final result, but rather to generate a novel hypothesis or confirm a hypothesis with different data.

We have shown that current tools don’t have the ability to incorporate clinical data with the omics data. One of the reasons might be that, to date, few clinical parameters are included when submitting omics data to the repositories. However, we postulate that just by having those parameters, more integrative analysis can be performed, allowing us to leverage the existing data in more depth. At the same time, our case studies associating pain and fatigue with their transcriptomics data show potential avenues of analysis, albeit some of the results did not yield many significant results.

Anyhow, we hope that due to recent developments in AI, more data will become available and RNAcare will allow users without a bioinformatics background to integrate data to generate new hypotheses.

## Availability and requirements

Project name: RNAcare

Project home page: https://github.com/sii-scRNA-Seq/RNAcare/

Operating system(s): Platform independent, tested on Linux (Ubuntu)

Compatible browsers: Firefox/Chrome

Programming language: Python, JavaScript

Other requirements: Python >= 3.8, Django >= 4.2, Nginx

## List of abbreviations

RNAcare: RNA-based Clinical Analysis and Research Engine
FAIR: Findable, Accessible, Interoperable, and Reusable
EDA: Exploratory Data Analysis
DGEA: Differential Gene Expression Analysis
DE: Differential Expression
IMID: Immune-mediated inflammatory diseases
PEAC: Pathobiology of Early Arthritis Cohort
ORBIT: Optimal management of rheumatoid arthritis patients who Require Biological Treatment
CPM: Count Per Million
VAS: Visual Analogue Scale
BMI: Body Mass Index
DAS28: Disease Activity Score using 28 joint counts
PCA: Principal Component Analysis
t-SNE: t-Distributed Stochastic Neighbour Embedding
UMAP: Uniform Manifold Approximation and Projection
REST: Representational State Transfer
CSR: Class Switch Recombination
ESR: Erythrocyte Sedimentation Rate

## Declarations

### Ethics approval and consent to participate

For the ORBIT data, all participants provided written, informed consent. The study protocol ORBIT [15] was approved by the West of Scotland Research Ethics Committee on 3/11/2009 (REC reference number: 09/S0703/109/ EudraCT number: 2009-011268-13)

For other studies, data were published, see RA-MAP [16] /PEAC [14].

### Consent for publication

We have consent for publication.

### Availability of data and materials

The code of can be found https://github.com/sii-scRNA-Seq/RNAcare/ and the webserver at https://rna-care.mvls.gla.ac.uk/

All data used are from published sources, as indicated in the Implementation section. The accession numbers are: E-MTAB-6141, GSE97810, GSE97948, GSE53986, GSE106382 and GSE20589.

### Competing interests

No competing interests.

### Funding

This work was funded through the IMID-Bio-UK consortium (MRC: MR/R014191/1). MT was funded through the IMID-Bio-UK project. WHH and TDO were supported by the Wellcome trust grant: 104111/Z/14/Z & A and TDO further by the ExposUM Institute of the University of Montpellier (grants ANR-21-EXES-0005 and Occitanie Region (TDO)).

### Authors’ contributions

MT designed and built RNAcare. He performed the data analysis, generated the figures and wrote the first draft. WHH gave technical support for implementation and tested the code. DP, SS, CG and FM discussed the clinical results. TDO did the conceptualisation of the project and wrote the manuscript. All authors contributed to the writing of the manuscript.

## Acknowledgements

We would like to thank Iain McInnes and the IMID-Bio-UK consortium for funding and making the data available. Further, we would like to thank Scott Arkison (School of Infection and Immunity) for maintaining the computational infrastructure that hosts RNAcare.

## Supplemental Material

**Supplemental Figure 1.**
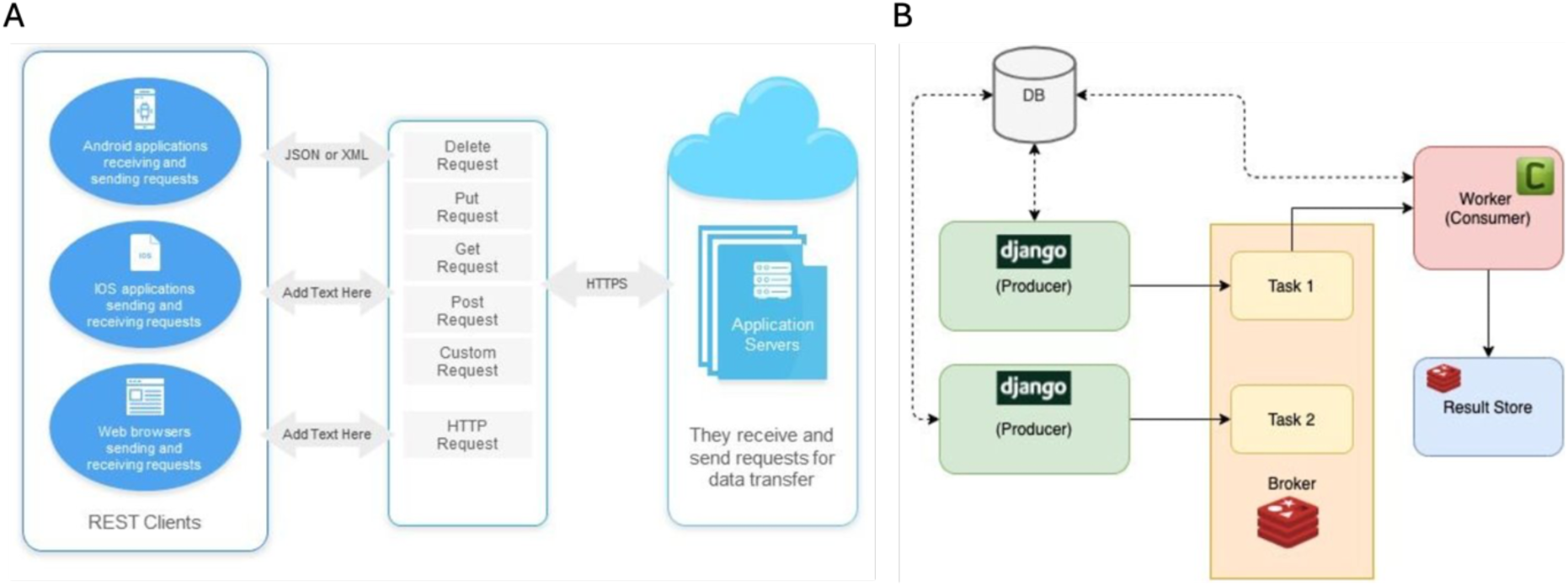
Platform Architecture. **(A)** Django is used to establish a RESTful webservice; **(B)** At the backend, Celery is used to establish a distributed scheduling system for multiple users.

**Supplemental Figure 2.**
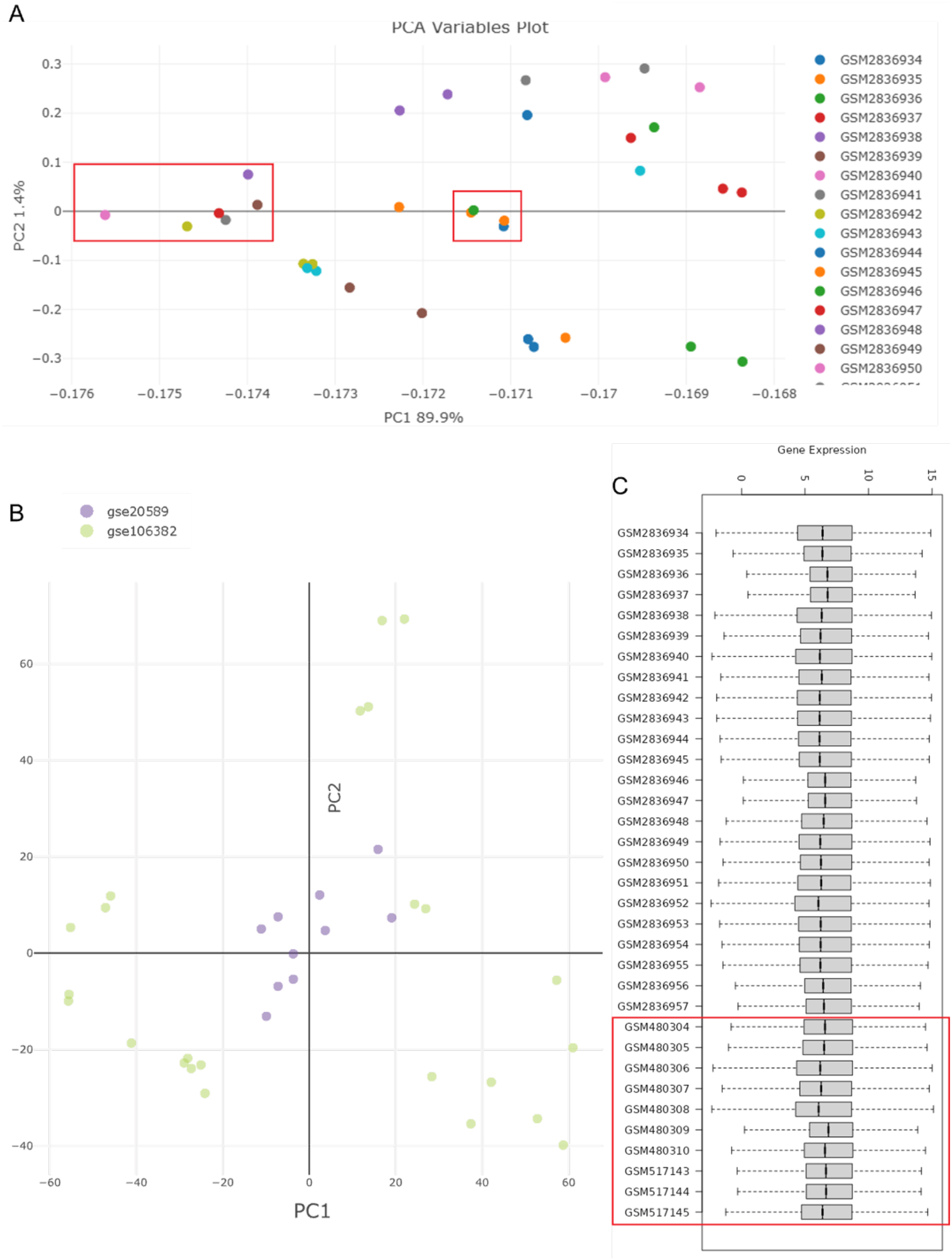
Comparison with GEOexplorer using GSE106382 and GSE20589. (A) PCA plot with GEOexplorer after removing batch effect. Points in the red boxes are GSE20589. (B) PCA plot by batches with RNAcare with log transformation after removing batch effect. We can see RNAcare provides a visualization by batches and can easily support combining more than 2 datasets. (C) Record comparison of GEOexplorer after batch correction.

**Supplemental Figure 3.**
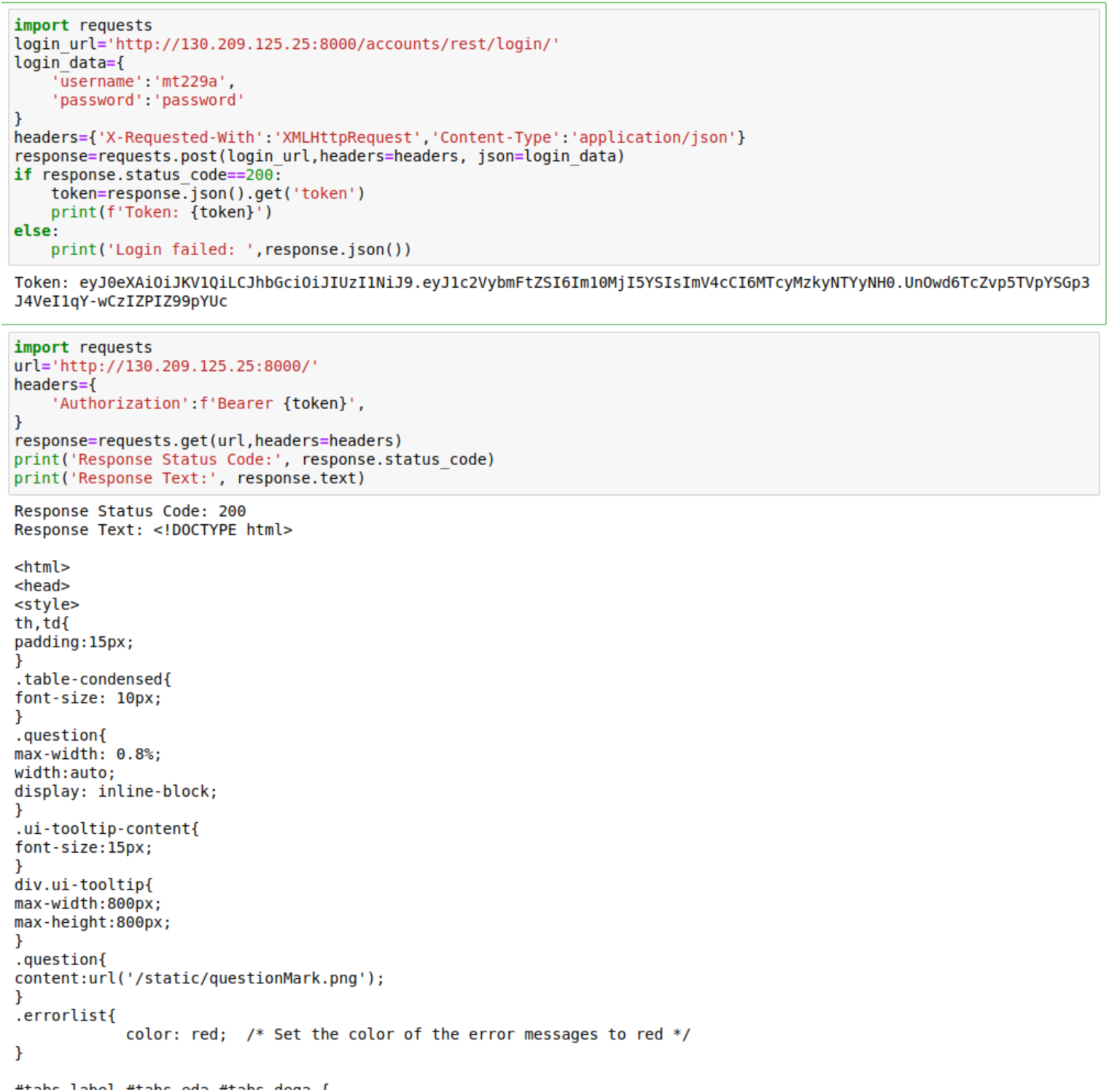
We simulate requests from clients, firstly, the client requests a token using an existing username and password, after getting the token, then the client can access the resources with the URLs

**Supplemental Figure 4.**
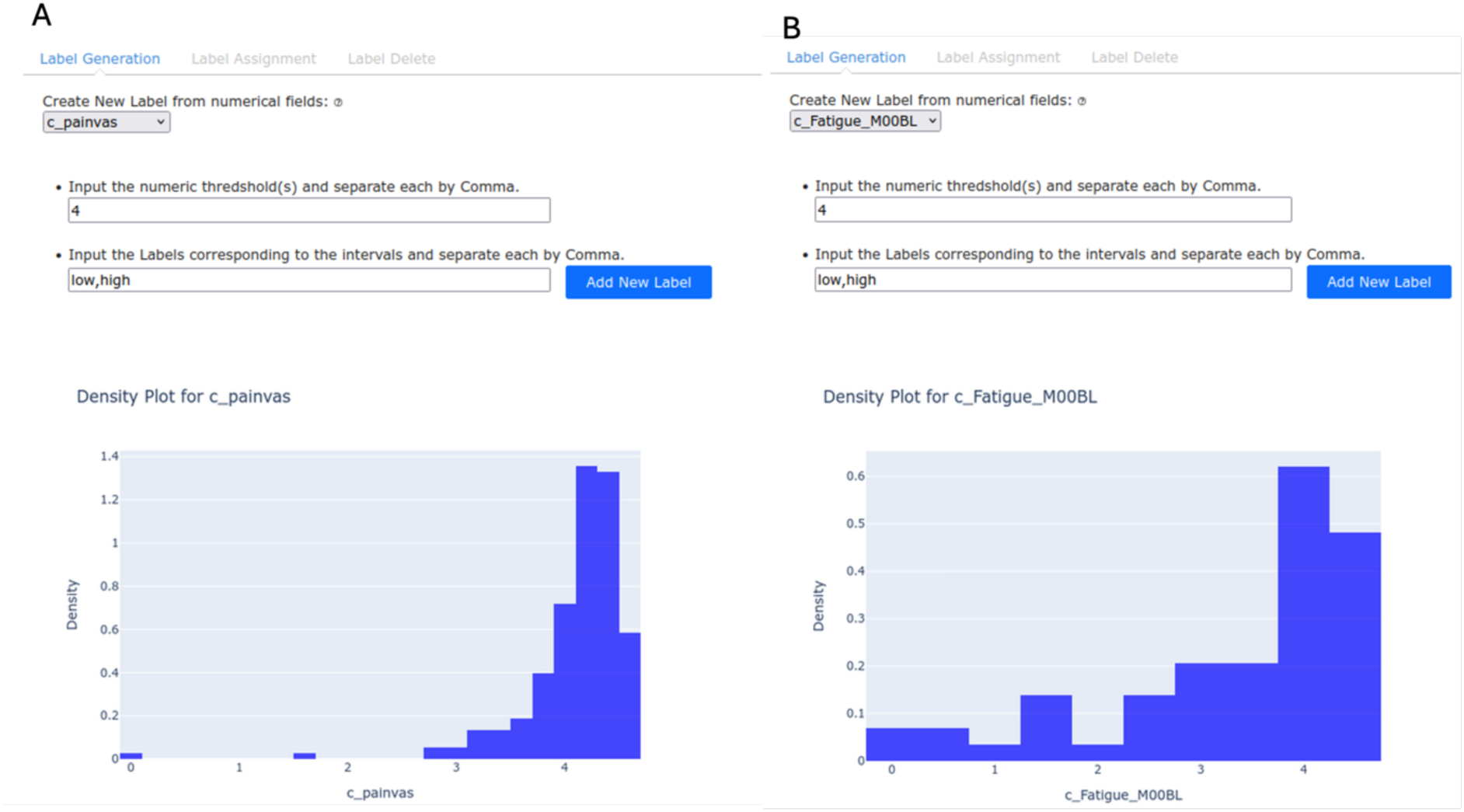
(A) Integration of PEAC and ORBIT with parameters settings. (B) Parameters settings for Fatigue (RA-MAP)

**Supplemental Table 1.** Comparison of different platforms. we consider existing software platforms for gene expression analysis. Although they have common characteristics, their implementations vary a lot, which affects their usability, scalability and robustness. For the comparison, we consider three aspects, same as what Kleverov et al. 2024 [9] did: (1) support for gene expression analysis steps, (2) data availability and (3) user experience. +/-* As processed in ARCHs4 and/or Dee2 projects.

| Application |  | GenePattern<br>Chapman et al.,2006[66] | GEO2R<br>Barrett et al., 2013[67] | BicOv-<br>erlapper<br>2 San-<br>tamaria<br>et al.,<br>2014[68] | Babe-<br>lomics<br>Alonso<br>et al.,<br>2015[69] | Mor-<br>pheus<br>Gould,<br>2016[70] | START<br>Nelson<br>et al.,<br>2017<br>[8] | BioJu-<br>pies<br>Torre et<br>al.,<br>2018[71] | iDEP Ge<br>et al.,<br>2018[6] | Degust<br>Powell,<br>2019 | GREIN<br>Mahi et<br>al.,<br>2019<br>[7] | EX-PANDER<br>Hait et al.<br>2019[72] | RaNa-<br>seq<br>Prieto<br>and Bar-<br>rios,<br>2019[73] | RNAde-<br>tector La<br>Ferlita et<br>al.,<br>2021[74] | GEOex-<br>plorer<br>Hunt et<br>al.,<br>2022[12] | Phanta-<br>sus.,<br>2024[9] | RNAcare |
| --- | --- | --- | --- | --- | --- | --- | --- | --- | --- | --- | --- | --- | --- | --- | --- | --- | --- |
| Gene<br>expres-<br>sion<br>analy-<br>sis key<br>steps | Normaliza-<br>tion | + | - | - | + | + | + | + | + | - | + | + | - | - | + | + | + |
|  | PCA | + | - | - | + | - | + | + | + | + | + | + | + | - | + | + | + |
|  | Clustering | + | - | - | + | + | - | + | + | - | - | + | - | - | + | + | + |
|  | Differential<br>Expression | + | + | + | + | - | + | + | + | + | + | + | + | + | + | + | + |
|  | Pathway<br>analysis | + | - | + | + | - | - | + | + | - | - | + | + | - | + | + | + |
|  | Batch inte-<br>gration | - | - | - | - | - | - | - | - | - | - | - | - | - | only two | - | + |
| Gene<br>expres-<br>sion<br>sources | user-pro-<br>vided data | + | - | + | + | + | + | + | + | + | - | + | + | + | - | + | + |
|  | GEO micro-<br>array | + | + | + | - | - | - | - | - | - | - | - | - | - | + | + | Upload Man-<br>ually/Auto-<br>mation under<br>development |
|  | GEO RNA-seq | - | - | - | - | - | - | Under develop-<br>ment | Under develop-<br>ment | - | + | - | + | - | - | Under develop-<br>ment | Upload Man-<br>ually/Auto-<br>mation under<br>development |
| User<br>experi-<br>ence<br>fea-<br>tures | Architecture | Web non-Shiny | Web non-Shiny | Local in-<br>stallation | Web non-Shiny | Web non-Shiny | Web Shiny | Web non-Shiny | Web Shiny | Web non-Shiny | Web Shiny | Local in-<br>stallation | Web non-Shiny | Local in-<br>stallation | Web Shiny | Web non-Shiny | Web Service/<br>Local instal-<br>lation |
|  | Saved ses-<br>sions | + | - | - | - | +/-* | - | + | - | - | - | - | + | - | - | + | - |
|  | Interactive<br>heatmap | - | - | +/-* | - | + | +/-* | +/-* | +/-* | + | + | + | +/-* | - | + | + | + |
|  | Interactive<br>plots | - | - | - | + | + | + | + | + | + | + | + | + | - | + | + | + |
|  | Editing<br>sample an-<br>notations | - | - | - | + | + | - | - | - | - | - | + | - | + | - | + | + |
|  | Editing gene<br>annotations | - | - | - | + | - | - | - | + | - | - | - | - | - | - | + | + |

**Supplemental Table 2.**
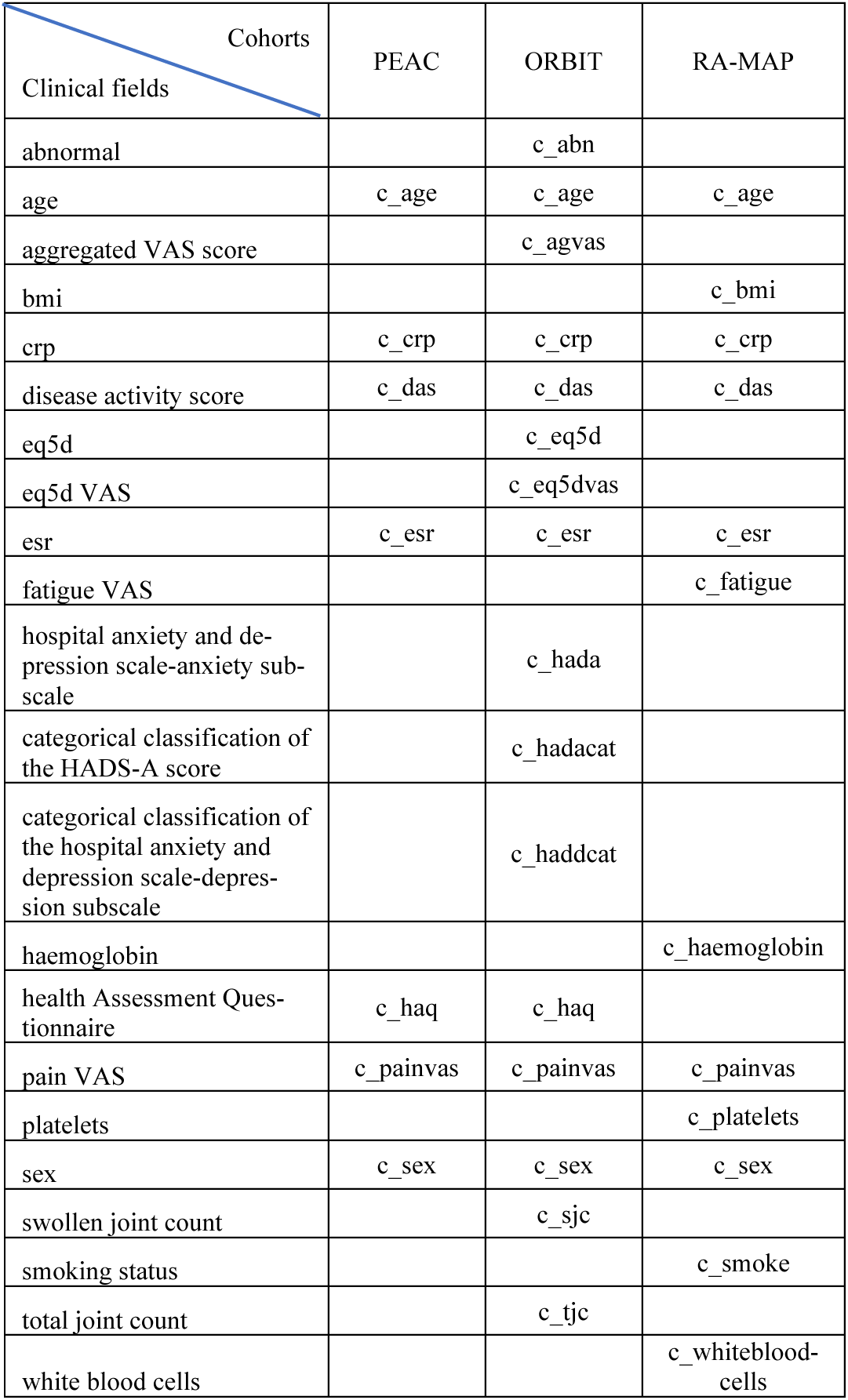
Clinical fields for datasets used in the paper.

## Notes

https://github.com/sii-scRNA-Seq/RNAcare/

### Competing Interest Statement

The authors have declared no competing interest.

